# The cryo-EM structure of the bacterial flagellum cap complex suggests a molecular mechanism for filament elongation

**DOI:** 10.1101/807677

**Authors:** Natalie S. Al-Otaibi, Aidan J. Taylor, Daniel P. Farrell, Svetomir B. Tzokov, Frank DiMaio, David J. Kelly, Julien R.C. Bergeron

## Abstract

The bacterial flagellum is a remarkable molecular motor, present at the surface of many bacteria, whose primary function is to allow motility through the rotation of a long filament protruding from the bacterial cell. A cap complex, consisting of an oligomeric assembly of the protein FliD, is localized at the tip of the flagellum, and is essential for filament assembly, as well as adherence to surfaces in some bacteria. However, the structure of the intact cap complex, and the molecular basis for its interaction with the filament, remains elusive. Here we report the cryo-EM structure of the *Campylobacter jejuni* cap complex. This structure reveals that FliD is pentameric, with the N-terminal region of the protomer forming an unexpected extensive set of contacts across several subunits, that contribute to FliD oligomerization. We also demonstrate that the native *C. jejuni* flagellum filament is 11-stranded and propose a molecular model for the filament-cap interaction.

## Introduction

The bacterial flagellum is a macromolecular motor that rotates and acts as a propeller in many bacteria. It is associated with virulence in many human pathogens including *Salmonella*, enteropathogenic *Escherichia coli, Campylobacter*, and *Helicobacter* species ^1,2^. The flagellum is composed of > 25 different proteins, and consists of three main regions: the basal body acts as an anchor in the bacterial membrane, and includes the apparatuses for rotation and protein secretion; the hook forms a junction which protrudes from the outer membrane; and the filament, consisting of multiple repeats of a single protein (flagellin), forms the propeller ^3^. The filament, that can be > 20 μm in length, is topped by a cap complex, that consists of several copies of the protein FliD. This complex initially attaches to the hook-filament junction, and using a yet unknown mechanism, assists in building the filament ^4^.

Low-resolution cryo-EM studies of the cap complex in *Salmonella enterica* have suggested that it consists of five copies of FliD (also known as HAP2), forming a “stool”-shaped complex with a core “head” domain and five flexible “leg” domains, that interact with the growing end of the filament ^5,6^. Crystal structures of the FliD head domain have been reported for several species, and revealed a range of crystallographic symmetries, from tetramers in *Serratia marscecens* (FliD_sm_), pentamers in *S. enterica* (FliD_se_) and hexamers in *E. coli* (FliD_ec_) and *Pseudomonas. aeruginosa* (FliD_pa_) ^7–10^. This observation led to the hypothesis that the cap complex can have a different, species-specific oligomeric states.

The flagellar filament has been studied extensively by cryo-EM, and its high-resolution structure has been reported in a range of bacteria, including *Bacillus subtilis, P. aeruginosa* and *S. enterica*. In all of these, the filament was shown to consist of 11 proto-filaments ^11,12^. However, a low-resolution cryo-EM study of the *C. jejuni* flagellar filament suggested the presence of 7 protofilaments ^13^. Taken together with the range of oligomeric states observed in the FliD crystal structures, these observations have led to a model where in different bacterial species, the cap complex has different oligomeric states (N), and in the corresponding filaments, the number of protofilaments is 2N + 1 ^7^.

*Campylobacter jejuni* is a Gram-negative, spiral-shaped microaerophilic epsilon proteobacterium, colonizing the lower gastrointestinal (GI) tract of humans and poultry ^14^. It is often the most common cause of bacterial gastroenteritis and can lead to severe sequelae such as Guillain-Barré (GBS) and Miller-Fisher syndromes (MFS) ^15^. *C. jejuni* has two polar flagella located at each cell pole, which have an important function not only in motility, but are also responsible for adherence to surfaces, and for the secretion of virulence factor proteins ^15,16^. FliD_cj_ is the major antigen in *C.jejuni* and thus a target for vaccine design ^17–19^.

In this study, we report the structure of the *C. jejuni* flagellar cap complex by cryo-EM. This structure demonstrates that FliD_cj_ is pentameric, with an extensive set of contacts across several residues at the termini, that contribute to stabilizing the oligomeric state. We show that these interactions are essential for cell motility. We also observe that the full-length FliD protein for both *S. marscecens* (FliD_sm_) and *P. aeruginosa* (FliD_pa_) also form pentamers, with similar dimensions to that of FliD_cj_, indicating that the pentameric state of FliD within the cap complex is likely universal. Finally, we demonstrate that the native *C. jejuni* flagellum filament is 11-stranded, similar to other known flagellum filament structures. These observations allow us to propose a molecular model for the filament-cap interaction, and cap-mediated filament elongation.

## Results

### Cryo-EM structure of the flagellum cap complex

Existing high-resolution structures of FliD have so far been limited to the head domain. We therefore sought to characterize the intact FliD protein. To that end, we purified full-length FliD from several species: *C. jejuni* (FliD_cj_), *P. aeruginosa* (FliD_pa_) and *S. marcescens* (FliD_sm_) (Figure S1a). Size-exclusion chromatography demonstrated that all three proteins form oligomeric assemblies (not shown). However, preliminary negative-stain analysis showed that while the complexes formed by FliD_pa_ and FliD_sm_ are heterogeneous (Figure S1b), FliD_cj_ forms homogeneous complexes, suitable for structural characterization.

Next we used cryo-EM to determine the structure of FliD_cj_. The protein forms discrete particles in vitreous ice, and 2D classification confirms that it adopts the dumbbell shaped structure previously reported for FliD_st_ (Figure S2a). In addition, a significant subset of particles adopted top-view orientations, with clear 5-fold symmetry. This allowed us to obtain a structure of the full complex, to 4.71 Å resolution (Figures S2b, S2e).

The FliD_cj_ complex possesses an overall architecture similar to FliD_sm_ ^5,6^, consisting of ten subunits, with two pentamers interacting in a “tail-to-tail” orientation, through the leg domains (Figure 1a). A pentamer is about 170 Å in height (the decamer is ∼300 Å) and 130 Å in width with a 20 Å lumen (Figure 1c). We note however that the map shows a wide range of local resolution, with the leg domain well defined and with visible side-chains, while the head domain is much more poorly defined (Figure S2b).

**Figure 1:**
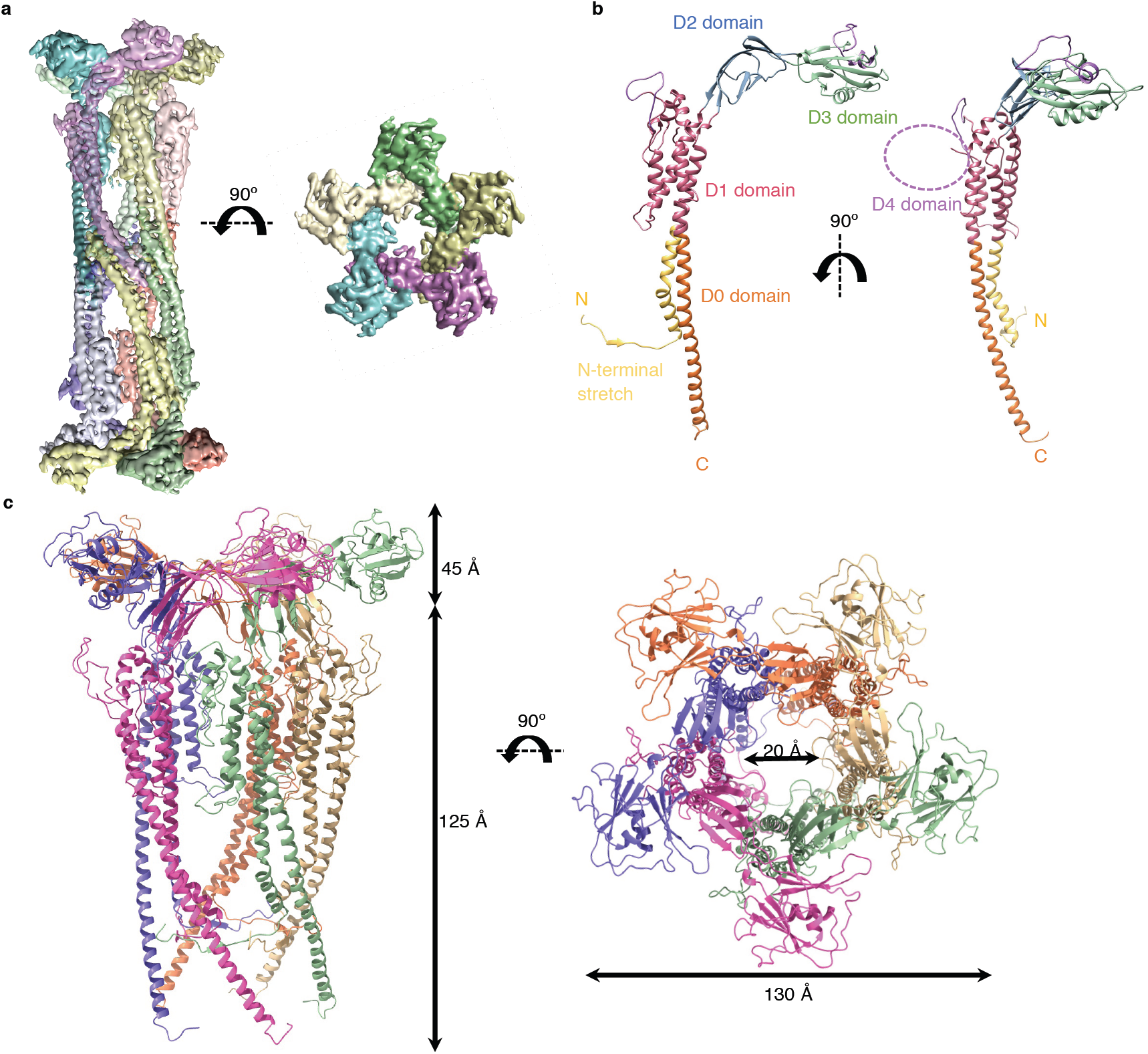
Structure of FliDcj. **(a)** The cryo-EM map of FliD_cj_, segmented and coloured by subunit. A side view of the full complex map is shown on the left, and a top view of the head domain map is shown on the right. **(b)** Cartoon representation of the full-length FliD_cj_ monomer, coloured according to domain organization. The purple dotted circle indicates the position of the D4 domain. **(c)** Cartoon representation of the FliD_cj_ pentamer, corresponding to the intact cap complex, with respective measurements. Side view (left) and top view (right) are shown, and color-coded as in (a).

This suggests that the complex is dynamic, with a hinge between the leg and head domains. To address this, we therefore performed a focused refinement on the head domain only, leading to a map at 5.02 Å resolution for this domain (Figure S2c, S2e). Using this map, we were able to generate an atomic model for this region of FliD_cj_, based on the crystal structure of FliD_ec_ (PDB ID: 5H5V) ^8^. We then used the map of the full complex to build the atomic model for the leg domain *de novo* (Figure S2d, S2f, Table 1).

**Table 1:** Maps and atomic model statistics.

|  | FliD<sub>Cj</sub> | Filament |
| --- | --- | --- |
| *Data collection* |  |  |
| Microscope | Titan Krios | Tecnai Arctica |
| Voltage (kV) | 300 | 200 |
| Camera | K2 Summit | Falcon III |
| Magnification | 36232 | 53000 |
| Pixel size (Å) | 1.38 | 2.03 |
| Defocus range (μm) | -1.0 to -2.6 | -0.8 to -2.0 |
| Total dose (eÅ<sup>-2</sup>) | 41 | 45 |
| Number of micrographs | 1223 | 100 |
| Total particles used | 55967 | 71828 |
| *Model Composition* |  |  |
| Non-hydrogen atoms | 37320 |  |
| Protein Residues | 4880 |  |
| *Refinement* |  |  |
| Resolution | 4.71 Å (full map)<br>5.02 Å (head domain) | 8.6 Å (symmetrical)<br>27.2 Å (asymmetrical) |
| Mask CC | 0.77 |  |
| Volume CC | 0.77 |  |
| *R.m.s. deviations* |  |  |
| Bond lengths (Å) | 0.005 |  |
| Bond angles (°) | 0.543 |  |
| *Validation* |  |  |
| MolProbity score | 1.85 |  |
| Clashscore | 8.36 |  |
| Poor rotamers (%) | 0.00 |  |
| *Ramachandran plot* |  |  |
| Favoured (%) | 94.13 |  |
| Allowed (%) | 5.87 |  |
| Disallowed (%) | 0.00 |  |

The FliD_cj_ structure shows that the FliD protomer folds in on itself in a ν-shape, which results in N and C termini next to each other in the leg domain. The overall architecture, as proposed previously, consists of a D0 domain formed by a long coiled coil, consisting of two helices located at the termini. A four-helix bundle forms the D1 domain. Connected to the D0-D1 leg domains are D2-D3 domains, rich in anti-parallel β-sheets, forming the head (Figure 1b) ^7–9^. This overall architecture is similar to that of the flagellin and hook, and in agreement with the previously reported structures of the FliD head domain ^20^. Intriguingly, while it was predicted that the D0 domain consists of a two-helix coiled-coil, as present in the flagellin and hook, our structure reveals that the N-terminal 17 residues are extended into a stretch that folds under and behind the monomer, interacting with the preceding subunit via a short β-strand. As a consequence, the C-terminal helix of the coiled-coil is not partnered with the N-terminus, but instead interacts with that of another molecule through hydrophobic interactions, forming the pentamer-to-pentamer interface. This intriguing architecture likely explains the strong tendency of FliD to form tail-to-tail complexes during isolation, as observed in FliD_se_ ^6^ and FliD_cj_ (this study).

We also note that FliD_cj_ possesses a long insert within the D1 helix bundle, not present in other orthologues (Figure S3a). Secondary structure prediction indicates that this insert is likely globular (not shown), leading to the hypothesis that it forms an additional domain, termed D4. This type of domain insertion is not unusual, and has been observed in other FliD orthologues, as well as in flagellin and hook proteins ^10,11,21,22^. In our FliD_cj_ map, we were able to observe density for this domain (Fig S3b), however it is at very low resolution, and did not allow us to build an atomic model. This suggests that the D4 domain is flexible. Indeed, further 3D classification revealed at least 4 distinct positions for this domain (Figure S3c). The role of this D4 domain is not known, but we postulate that it could be related to FliD_cj_’s capacity to bind to heparin, a feature involved in *C. jejuni* adherence but not observed in other FliD orthologues^4^.

### Comparison with other FliD orthologues

Our cryo-EM structure of FliD_cj_ is the first high-resolution structure of an intact FliD protein. Nonetheless the crystal structure of the head domain, corresponding to domains D2-D3, has been reported for a range of species, including *S. enterica, P. aeruginosa, E. coli, S. marcescens*, and *H. pylori* ^6–10^. In all orthologues, the structure is very similar, with RMSB values ranging from 1.5 Å to 2.5 Å to that of FliD_cj_ (Figure 2a). In the *E. coli* orthologue, domain D1 was also present in the structure. It consists of a 4-helix bundle, and this structure is very similar to that of FliD_cj_, with a RMSD of 1.5 Å between the two structures. Nonetheless, we note that the position of D1 relative to that of D2-D3 is dramatically different in FliD_ec_ compared to FliD_cj_ (Figure S4a). This suggests that the hinge between D1 and D2 is flexible, as supported by our focused refinement result.

**Figure 2:**
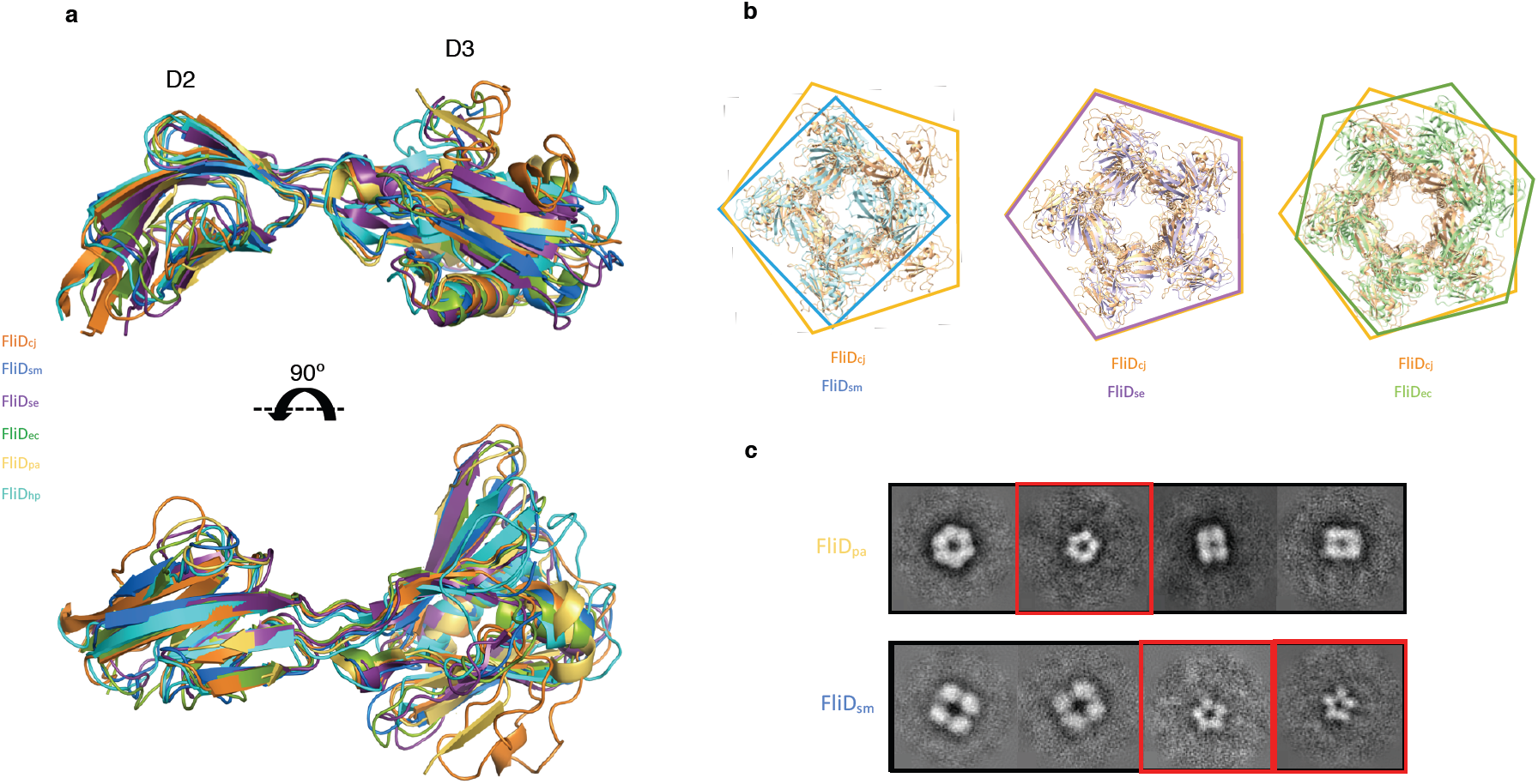
Comparison of the FliD structures across bacterial species. **(a)** Overlay of the FliD D2-D3 domains structures from *C. ejuni* (FliD_cj_, 6SIH – this study, orange), *S. marcescens* (FliD_sm_, 5XLJ, blue), *S. enterica* (FliD_se_, 5H5T, purple), *E. coli* (FliD_ec_, 5H5V, green), *P. aeruginosa* (FliD_pa_, 5FH5, yellow), and *H. pylori* (FliD_hp_, 6IWY, cyan). **(b)** Alignments of X-ray crystallography derived oligomeric structures of the head domains from FliD_sm_, FliD_se_ and FliD_ec_, to that of the FliD_cj_ pentamer. The diameter of the lumen as well as the outer diameter of the capping protein is similar in the pentamer formed by FliD_se_, but significantly smaller in the tetramer formed by FliD_sm_, and larger in the hexamer formed by FliD_ec_. **(c)** 2D classes obtained from negative stain data of recombinant FliD_pa_ and FliD_sm_ Figure S1b). FliD_pa_ shows 3 different sets of particles: a class with particles showing 6-fold symmetry, a class showing smaller particles with 5-fold symmetry, and classes of particles with 4-fold symmetry. FliD_sm_ shows 2 different sets of particles: large particles with 4-fold ymmetry, and smaller particles with 5-fold symmetry. The size of the 5-fold symmetry particles in both orthologues is similar to that observed in 2D classification of FliD_cj_. The red squares indicate classes with particles of similar shape and dimensions to the FliD_cj_ tructure, with distinctive five-fold symmetry.

In our cryo-EM map, FliD forms a pentameric architecture, consistent with the low-resolution cryo-EM structure of FliD_se_, with a similar overall architecture consisting of two pentamers in a head-to-tail arrangement. In contrast, crystal structures of the head domains from FliD in several species reported a range of oligomeric states, including tetramer (FliD_sm_), pentamer (FliD_se_) and hexamers (FliD_pa_ and FliD_ec_)^7–9^. When comparing the dimensions of these structures, the diameters of all complexes are similar, around ∼140 Å. However, the dimension of the lumen differs significantly between structures, with FliD_cj_ and FliD_se_ having a central lumen of ∼20 Å, while FliD_pa_ and FliD_ec_ have a lumen of ∼50 Å and ∼40 Å respectively, and FliD_sm_ a ∼15 Å lumen (Figure 2b). Even in the case of FliD_se_, which crystallized as a pentamer, while the overall dimensions are similar to that of the head domains of the FliD_cj_ pentamer, in the *E. coli* orthologue the pentamer is flattened compared to that of FliD_cj_ (Figure S4b). Based on our structure, we hypothesize that there is a large degree of plasticity in the interface between the D2-D3 domains of adjacent molecules, and therefore in the absence of D0, a range of interfaces can be trapped in the crystal contacts. We propose that the additional contacts formed by the N-terminal stretch are essential for FliD to adopt its true oligomeric state.

To verify this, we investigated the oligomeric state of full-length FliD_sm_ and FliD_pa_, the head domains of which crystallized as tetramers and hexamers, respectively, by negative stain TEM. As mentioned above, these proteins do not form uniform complexes (Figure S1b). Nonetheless, we noted that the majority of the particles appeared as top views, which allowed us to perform preliminary 2D classification to determine their lateral symmetry. This revealed that both orthologues form pentamers with similar dimensions to that of FliD_cj_ (Figure 2c). However, in the FliD_pa_ sample we observed additional particles with 6-fold and 4-fold symmetry, while in the FliD_sm_ sample there was a large percentage of particles with 4-fold symmetry. The dimensions of the particles in those 2D classes are significantly larger than the FliD pentamer, and therefore we could not conclude if these correspond to alternative oligomeric species, or to other negative stain artifacts and/or non-specific aggregates. However, the presence of pentamers with similar dimensions to that of FliD supports the hypothesis that the native architecture of the cap complex is a FliD pentamer, with contacts at the N-terminus required for FliD to adopt its true oligomeric state.

### Hydrophobic interactions in the D0 domain are required for forming functional filaments

As mentioned above, our structural characterization of the cap complex indicates an unusual architecture of the N-terminus, which forms a stretch that wraps around and forms contacts with two adjacent subunits, through hydrophobic contacts (Figure 3a). In particular, Residues Leu 9 and Phe 11 are buried within a pocket formed by Trp 614 and Tyr 617, located in the C-terminus of the adjacent molecule. This is of particular interest since it was shown that the C-terminus contributes to the oligomerization of FliD and interaction with its chaperone ^23^. We also note that both the N- and C-termini of FliD are highly conserved across species, with mainly aromatic side-chains present in all orthologues in the aforementioned positions.

**Figure 3:**
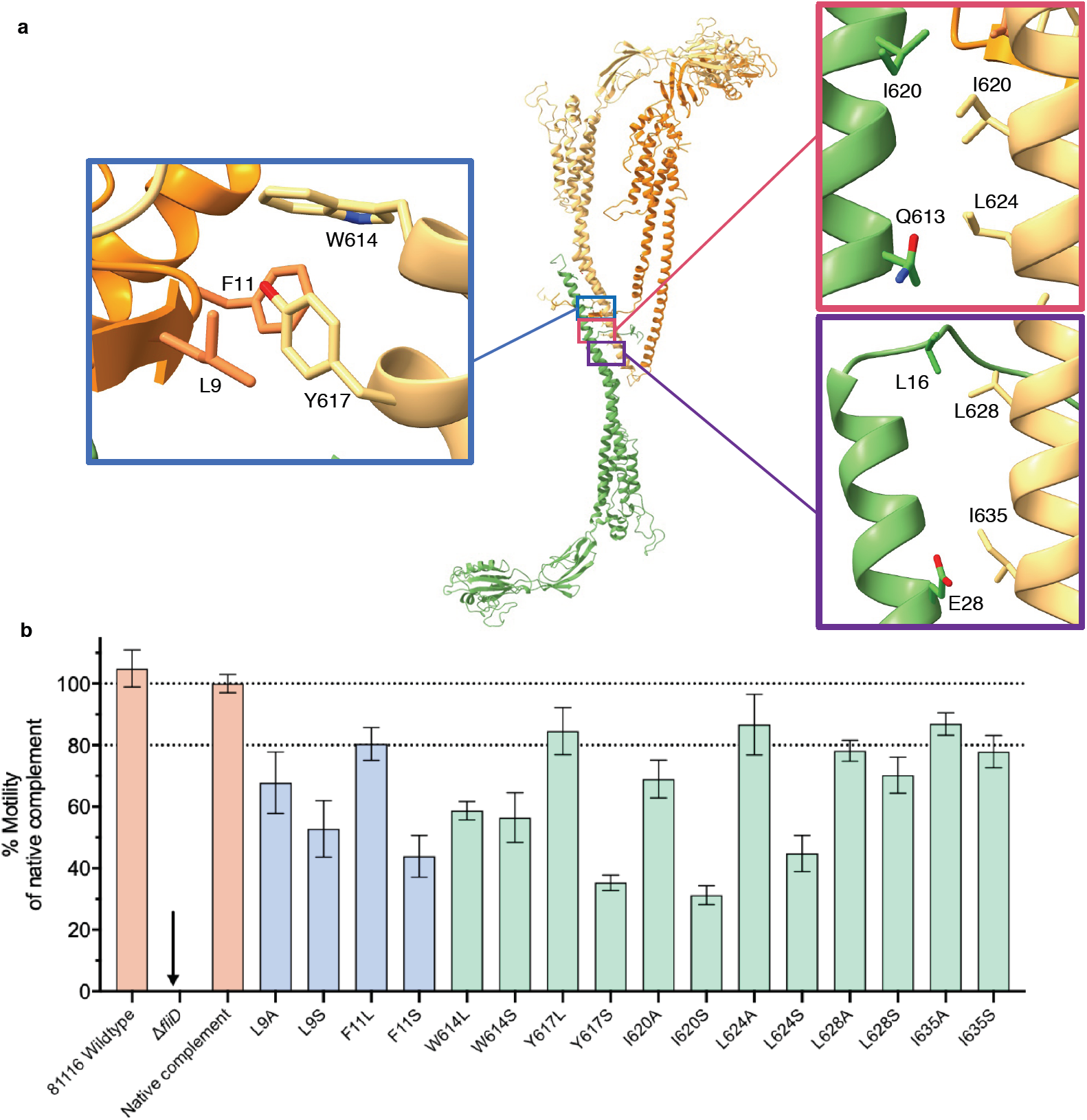
FliDcj amino-acid substitutions and their effect on motility. **(a)** Location of the mutated residues on the FliDcj structure. Adjacent subunits (Yellow, Orange) were chosen to represent the interaction of the N-terminus residues with that of the next subunit C-terminus and which might be important in the formation of the pentamer. The residues in the lower C-terminus were chosen to represent the interaction with flagellin upon flagellar elongation, as the conserved residues in that region which bind to the bottom subunit C-terminus might mimic the interactions with the flagellin monomer. **(b)** Motility assay results for point mutants represented as the mean percentage of the native complement strain, based on swarm diameter on soft agar. Controls (WT, deletion mutant and complement) are in orange. C-terminal mutants in light green and N-terminal mutants in purple. Error bars show standard deviation.

To confirm the role of these residues in FliD function, we engineered a *C. jejuni fliD* knockout strain (Δ*fliD*), leading to a loss of motility in a soft agar swarm assay. Accordingly, no filament was observed in this strain (Figure S5a). Genetic complementation by expressing the *fliD* gene at a distal site on the chromosome fully rescued motility (Figure 3b, S5a), and we exploited this to engineer point mutations in the aforementioned residues to assess their impact on motility.

Mutation of Leu 9, Phe 11, Trp 614 or Trp 617 significantly reduced motility, up to 40% for mutations to polar residues (F11S, W614S, L9S or Y617S) (Figure 3b). This confirms that the hydrophobic properties of these residues are critical for motility, suggesting that the interaction formed by the N-terminal stretch contributes to FliD function. To verify if motility was affected because the aforementioned mutations prevented filament assembly, we visualized the corresponding bacteria by TEM. All of the mutations still led to bacteria with assembled filaments, of length similar to that of WT bacteria (Figure S5b), demonstrating that the corresponding FliD proteins are still able to promote filament elongation. However, we noted that the filaments are much more brittle in the mutants, with between 60 and 80% of filaments found unattached to the bacterial cell, versus ∼ 20% in the WT bacteria (Figure S5c). We also note that the N-terminal ∼ 20 residue stretch corresponds to the secretion signal in flagellar filaments of *S. enterica,* so potentially a similar signal exists for FliD to be secreted through the flagellum T3SS ^24^. The observation that in the mutants described above, the filament is still formed, is a strong confirmation that these mutations did not interfere with FliD secretion, but rather with its function to promote filament elongation. The second set of interactions observed in the D0 domain, is formed between the C-terminus of FliD in the pentamer-to-pentamer interface (Figure 3a). Evidence from tomography, as well as other biochemical data, indicate that this interaction is not physiological ^5,25–27^. However, since it is observed in both FliD_cj_ and FliD_se_, we postulated that it mimics the interaction between FliD and the filament. To verify this, we engineered a series of mutations in the residues forming this interface (Leu 628, Ile 635, Leu 624 and Ile 620) and characterized their impact on motility as described above (Figure 3b). Mutating these residues impacted motility, however the effect is less pronounced than the mutants involved in the N-terminal stretch interaction, with the exception of I620S and L624S mutations. We propose that this is because the overall hydrophobic propensity, rather than specific contribution of each amino acid, is the critical element of this region of the protein.

### A structural model of the *C. jejuni* filament

Current Cryo-EM structures of various flagellar filaments have demonstrated that they consist of 11 protofilaments, formed by a single protein, the flagellin ^11^. The flagellin consists of four domains D0-D3, and can adopt two conformations, termed L and R, leading to two alternative filament structures, left-handed and right-handed, respectively. *C. jejuni* possesses two flagellin homologues, FlaA and FlaB, that are ∼ 95% identical to each other, with both required for the formation of fully functional filaments. FlaA and FlaB are highly similar to other flagellins (Figure S6a), except for an ∼ 70 amino acid insert in D2 that likely consists of a globular insert, as observed in several flagellin orthologues ^11^. Surprisingly, a previously published EM structure of the *C.jejuni* filament had reported a 7 protofilament arrangement ^13^. However, this structure was obtained from a FlaA G508A mutant, in the absence of FlaB, and is at low resolution. It is therefore not clear if this was an artifact and/or wrong interpretation of the data, or if the *C. jejuni* filament indeed possesses a different architecture to other species.

To reconcile this, we sought to determine the structure of the native *C. jejuni* filament, directly from wild type cells (Figure 4a). To avoid biases due to symmetry, we initially performed a reconstruction without any helical symmetry applied. This map clearly possessed 11-fold symmetry (Figure S6b), despite the low resolution (∼ 27 Å, figure S6c). This demonstrates that the *C. jejuni* flagellar filament consists of 11 protofilaments with a lumen of ∼25-30 Å and outer diameter of ∼200 Å, similar to that of other bacterial species. We therefore refined the map further by applying helical symmetry, with a 65.4° twist and 7.25 Å rise, which allowed us to reach ∼ 8.6 Å resolution (Figure S6c). In this map the central D0-D1 domains are well resolved, with the density for helices clearly visible (Figure 4b). The density for domains D2 and D3 is visible, but less well resolved. The fact that we can only reach limited resolution is perhaps not surprising, since we likely have a combination of L and R conformations for the flagellin.

**Figure 4:**
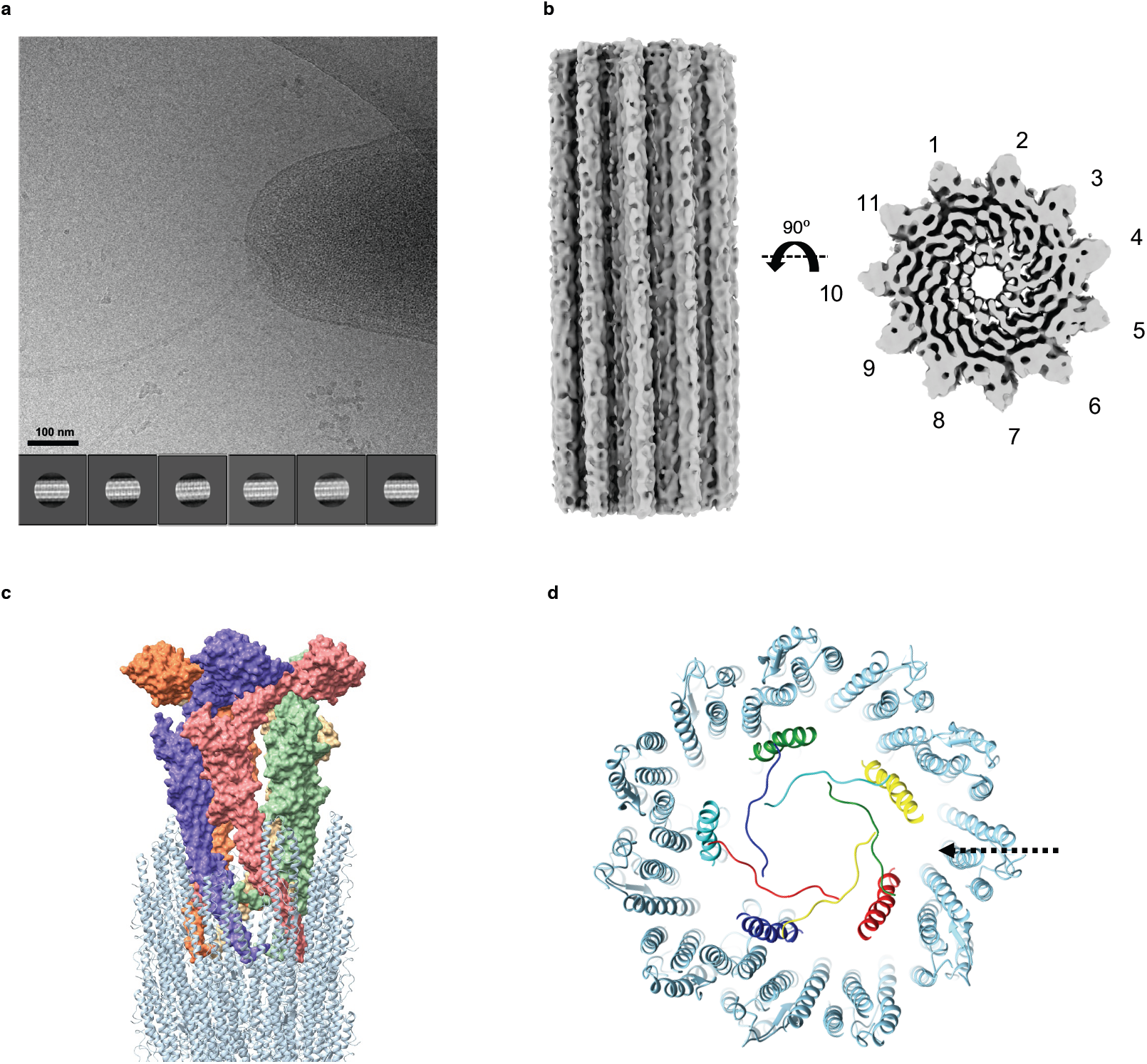
Structural characterization of the native *C. jejuni* flagella. **(a)** Cryo-electron micrograph of the native *C. jejuni* flagella, used for the 3D reconstruction. 2D classes, generated from ∼71828 particles, are shown below. **(b)** EM map of the native flagellum, with helical symmetry applied, to 8.6 Å resolution. A side view is shown to the left, and top view to the right. **(c)** Structural model of the filament-cap complex, with FliD in surface representation and coloured as in Figure 1, and the filament atomic model in cyan. **(d)** Closeup view of the various FliD-flagellin interfaces in the model shown in (c). Both termini, with the N-terminal stretch and the C-terminal helix, are shown for each FliD protomer.

Based on this, we used the previously published structure of the *P. aeruginosa* filament (PDB ID: 5WK6), to position the cap complex within the filament structure. This allowed us to propose a model for FliD-flagellin interaction (Figure 4c). In this model, the C-terminus of FliD forms broadly non-specific, hydrophobic contacts with exposed regions of the filament, similar to flagellin-flagellin interactions (Figures 4c and 4d). A gap between adjacent FliD molecules, on the side of the leg domain, is positioned in a suitable location for the insertion of a flagellin molecule and is the likely site of exit for nascent molecules (Figure 4d). This however remains to be verified experimentally.

## Discussion

In previous studies, evidence suggesting different stoichiometries for the flagellum filament and/or cap complex in different species was based on low-resolution cryo-EM structures, and crystallographic symmetries of truncated proteins. Here we largely resolve this conflicting evidence, by demonstrating that FliD adopts a pentameric stoichiometry in a range of species, and that the filament of *C. jejuni* is 11-stranded, and not 7-stranded as reported previously. We can therefore conclude that the stoichiometry of these proteins is conserved across species, with a 11-to-5 asymmetry between these two different regions of the bacterial flagellum. Our structure of the intact cap complex, supported by mutagenesis studies, suggests that the FliD C-terminal domain interaction with the opposite pentamer in the decametric complex mimics that of the FliD interaction with the filament. We hypothesize that exposed hydrophobic residues, both on the D0 domain of flagellin molecules and in the C-terminus of FliD, act as a chaperonin-like environment to promote the folding and insertion of new flagellins ^28,29^.

Based on the results reported in this study, we propose a universal mechanism for cap-mediated filament elongation, as illustrated in figure 5. The FliD cap pentamer fits into the flagellum filament, through interactions between D0 of flagellins and the C-terminus of FliD. Because of the symmetry mismatch, this interaction is not present on one side of the cap complex. New flagellin molecules are secreted through the filament, and ultimately enter a chamber inside the cap complex (1). The flagellin then exits this cavity through a lateral opening, where the location of the next flagellin insertion site is positioned (2). The four other cavities are sterically blocked by the flagellum filament. There, exposed hydrophobic residues act as a chaperone, and promote flagellin folding in its insertion site (3). The folding of the new flagellin protomer leads to dislodging of the cap complex, that rotates by ∼ 35 ° (4), thus positioning an adjacent cavity of the cap complex close to the next flagellin insertion site (Figure 5).

**Figure 5:**
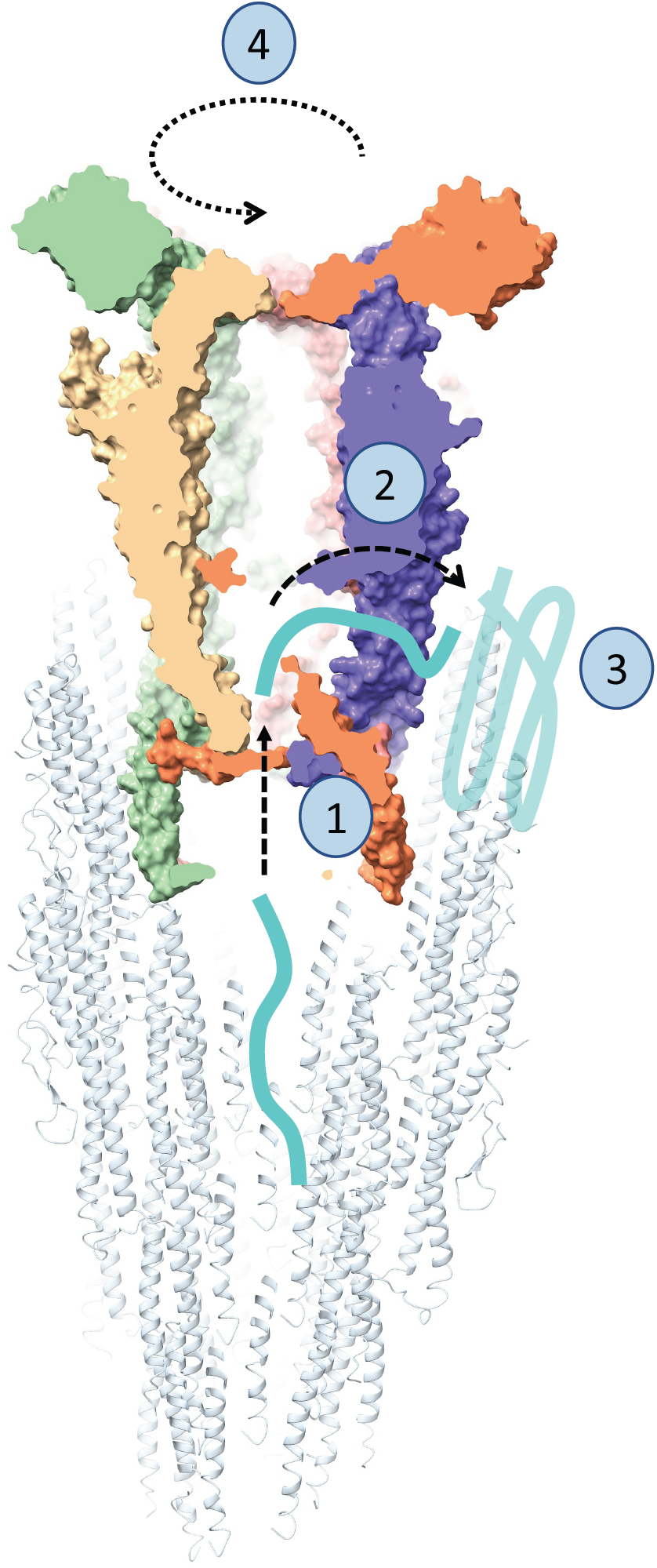
Model of cap-mediated filament elongation. **Nascent f**lagellin molecules are secreted through the filament, and enter the cap complex chamber (1). They then exit the cap complex through a side cavity (2), which positions them in close proximity to the site of insertion. There, hydrophobic patches, composed of both exposed flagellin molecules, as well as FliD D0 domain, act as chaperones, and promote flagellin folding (3). Following the insertion of a new flagellin subunit, the cap complex rotates (4), positioning it to have the open cavity towards the next site of insertion.

We note that previous studies, based on low-resolution tomography data, have suggested that the D0 domain of FliD might be dynamic, with the leg domains opening and closing to promote filament elongation ^5,9,26^. Our structure of the cap complex does not support this model, as we show that the N-terminal stretch of FliD is essential for filament elongation and maintains the leg domains in a rigid position. Our data supports an alternative mechanism, which had been proposed previously, whereby the cap complex acts as a rigid cog that rotates during flagellum elongation ^8^. Further experiments to characterize the flagellum-cap complex at high resolution will be required to confirm this model.

In conclusion, we report the cryo-EM structure of the flagellum cap complex, and demonstrate that FliD across multiple species (FliD_sm_, FliD_pa_ and FliD_se_) forms pentameric complexes. We show that the interface between opposite D0 leg domains in the FliD decamer complex is essential for cell motility and formation of a functional filament, and therefore likely plays a role in FliD-filament interactions. We also demonstrate that the *C. jejuni* flagellar filament possesses the same architecture as that of other species. Taken together, these results allow us to propose a universal model for cap-filament interaction as well as propose a mechanism for cap-mediated filament elongation.

## Materials and Methods

### Protein Expression and purification

The genes coding for FliD_cj_, FliD_sm_ and FliD_pa_, codon-optimized for expression in *E. coli*, were synthesized (BioBasic) and sub-cloned into pET28a (Novagen). Recombinant proteins were expressed in *E. coli* BL21-CodonPlus(DE3)-RIL cells containing the corresponding plasmids. For FliD_cj_, transformants were grown in LB medium at 37 °C until they reached log phase, and expression was induced by the addition of 1mM IPTG overnight at 20 °C. For both FliD_pa_ and FliD_sm_, expression was auto-induced in ZYM-5052^30^ media at 20 °C overnight. For all three proteins, cells were collected by centrifugation, resuspended in 50 mM HEPES 150 mM NaCl pH 7 and sonicated. The lysate was centrifuged at 14 000g at 4 °C for 45 minutes. The supernatants were applied onto a 5 ml HisPure™ Ni-NTA resin (ThermoScientific) gravity-based column equilibrated with 50 mM HEPES 150 mM NaCl pH 7 and eluted using a linear 20-500 mM Imidazole gradient. Fractions containing FliD were pooled and applied to a HiLoad Superdex 200 16/600 column (GE Healthcare) equilibrated with 50 mM HEPES 150 mM NaCl pH 7 for FliD_cj_ and 50 mM Tris 150 mM NaCl pH 8 for FliD_pa_ and FliD_sm_.

### Negative-stain grid preparation and data collection

For negative-stain TEM experiments, ∼ 5 μl of purified protein, or of cell culture in log phase, was applied onto glow-discharged, carbon-coated copper grids (Agar Scientific). After incubating the sample for ∼2 minutes at room temperature, the grids were rapidly washed in three successive drops of deionized water and then exposed to three successive drops of 0.75% uranyl formate solution. Images were recorded on a CM100 TEM (Phillips) equipped with a MSC 794 camera (Gatan) (FliD_cj_ and *C. jejuni* cell cultures) or a Technai T12 Spirit TEM (Thermo Fisher) equipped with an Orius SC-1000 camera (Gatan). Datasets were manually acquired with a pixel size of 2.46 Å/pix, and a defocus range from −0.8 μm to −2.0 μm. The micrographs were processed using cisTEM ^31^ package, with CTF parameters determined by CTFFIND4 ^32^. Approximately 3500 particles were picked for FliDsm and 2700 for FliDpa to generate representative two-dimensional (2D) class averages with 330 Å mask diameter. The point mutant flagella attachment was determined through imaging grids at 700x magnification at about 20 micrographs per mutant containing cell count from 30 to 100 cells. The percentage of attachment was calculated as a proportion of the total flagella observed per mutant.

### Cryo-EM grid preparation and data collection

For the structural characterization of FliD_cj_, aliquots of (5 μl) of purified protein at a concentration of 1 mg ml^-1^ was deposited onto glow-discharged C-flat holey carbon films 1.2/1.3 200 mesh (EMS). A Vitrobot Mark III (FEI) plunge-freezing device was used for freeze-plunging, using double-blotting ^33^ with a final blotting time of 6.5 seconds. Cryo-EM data were collected with a Titan Krios TEM operated at 300 kV and equipped with an energy filter (Gatan GIF Quantum) and recorded on a K2 Summit direct electron detector (Gatan) operated in counting mode. 1223 micrographs were automatically acquired with the EPU software (Thermo Fisher), at a pixel size of 1.38 /pix, using a total dose of 41 e^-^ Å^-2^ and with 40 frames per micrograph. The defocus range used for data collection was −1.0 μm to −2.6 μm.

For the structural characterization of the native *C. jejuni* filament, wild-type *81116* strain cell culture grown to OD_600_ = 5 was applied onto glow-discharged C-flat holey carbon films 2/2 200 mesh (EMS). A Leica EM GP (Leica) plunge-freezing device was used for freezing, with a 6 s blotting time. Cryo-EM data were collected on a Technai Arctica TEM (Thermo Fisher) operated at 200 kV and equipped with a Falcon III camera. 100 micrographs were collected using the EPU software (Thermo Fisher) in linear mode, with a pixel size of 2.03 Å/pix, with a total dose of 45 e^-^ Å^-2^ and 1 frame per micrograph. The defocus range used for data collection was approximately −0.8μm to −2.0 μm.

### Cryo-EM image processing and reconstruction

For FliD_cj_, processing was done in RELION 2.0 ^34^. Motion correction was performed with MotionCor2 ^35^, with dose-weighting. CTF parameters were determined by CTFFIND4 ^32^ software. Approximately 2000 particles were manually picked from selected micrographs to generate representative 2D class averages. These classes were used as templates for automated particle picking for the entire dataset. A total of 130000 particles were picked and extracted using a 280 × 280 pixels box. After multiple rounds of 2D classification, 55967 particles from the best 2D classes were obtained and used to generate an initial model. Following further 3D classification and refinement with D5 symmetry, a final map to 4.71 Å resolution was generated, which was sharpened using PHENIX 1.13 ^36^. The leg domains were visibly at a higher resolution than the head domains, therefore a mask centering on the head domain was used for further refinement with C5 symmetry, leading to a map of the head domain to 5.02 Å resolution. Further 3D classification of the masked head domain was used to identify 4 different conformations of the D4 domain not resolved in the full map. For the native *C. jejuni* filament, processing was done in RELION 3.0 ^34^. Motion correction was performed with MotionCor2 ^35^, with dose-weighting. CTF parameters were determined by CTFFIND4 ^32^. Filaments were manually picked, and particles were extracted using a 7.6 Å rise and 300 pixel box leading to a set of 254041 segments. Multiple rounds of 2D classification gave a final dataset of 71828 good particles which were used for 3D refinement, both with and without imposed helical symmetry. Without symmetry, the structure refined to 27.2 Å resolution, but when helical symmetry was applied, the final resolution after further classification and refinement was 8.6 Å, with a 65.4° twist and a 7.25 Å rise.

### Model building and refinement

For the D2-D3 domains, a homology model was generated with PHYRE2 ^37^, using the FliD_ec_ crystal structure ^8^ (PDB:5H5V) as a template. These domains were fitted into the sharpened map in Chimera ^38^. This model was subjected to iterative rounds of real-space refinement and building using PHENIX 1.16 ^36^ and Coot^39^ respectively. The N-terminal stretch was modeled with RosettaES ^40^, and then the remaining missing loops were modeled using RosettaCM^41^ guided by the electron density. The output model was refined once more in Coot to improve the geometry and delete any modelled residues in areas without electron density.

### Cultivation of *C. jejuni*

*C. jejuni* strain 81116 was grown on blood agar plates (Colombia base agar with 5% v/v defibrinated horse blood) in a microaerobic cabinet (Don Whitley, UK) at 42°C with a controlled atmosphere of 10% v/v O_2_, 5% v/v CO_2_ and 85% v/v N_2_. Where appropriate, the selective antibiotics kanamycin and chloramphenicol were added at 50 µg/ml and 20 µg/ml, respectively.

### Construction of *fliD* deletion mutant and complemented strains

A *fliD* mutation vector was constructed using NEB HiFi DNA assembly method (E2621, New England Biolabs). Briefly, flanking regions of *fliD* were amplified from *C. jejuni* 81116 genomic DNA using primers fliDmutantF1-R2 (Table S1). These flanks were assembled into pGEM3ZF either side of a non-polar kanamycin resistance cassette, amplified from pJMK30 using primers KanF/R (Table S1). The final mutation vector was designed such that spontaneous double crossover with the *C. jejuni* 81116 genome would result in the replacement of the majority of the open reading frame of *fliD* with the kanamycin resistance cassette, allowing a means of selection. For complementation of the mutant, *fliD* was amplified from *C. jejuni* 81116 genomic DNA using primers fliDcompF/R (Table S1). The amplified fragment was digested with BsmBI at sites incorporated into the primers and ligated into similarly digested pCmetK plasmid, a complementation vector for *C. jejuni* incorporating flanking regions of the pseudo-gene region corresponding to *cj0046* in *C. jejuni* 11168 to allow insertion into the genome, a constitutive promoter from the *C. jejuni metK* gene to drive expression of *fliD*, and a chloramphenicol resistance cassette. To generate the strains, wildtype *C. jejuni* 81116 was first transformed with the *fliD* mutation vector by electroporation and colonies selected for kanamycin resistance on blood agar plates. The isolated mutant strain was then further transformed with the *fliD* complementation vector and selected for double kanamycin / chloramphenicol resistance.

### Construction of *fliD* point mutants in *C.jejuni*

Point mutations in *fliD* were constructed by site directed mutagenesis of the complementation vector using the KLD method (M0554, New England Biolabs). Briefly, the *fliD* complementation plasmid was amplified by PCR with divergent primers containing targeted nucleotide substitutions in the forward primer (listed in Table S2). An aliquot of the linear PCR product was treated with the KLD enzyme mix to circularise the mutated plasmid while degrading any residual template. The treated plasmids were transformed into *E. coli* DH5α and transformants selected by chloramphenicol resistance. Plasmid was purified from multiple transformants and the *fliD* open reading frame was sequenced to ensure the correct substitution had been introduced without secondary mutations (LightRun sequencing, Eurofins EU). Point mutated complementation vectors were then transformed into the *C. jejuni fliD* mutant strain as above to generate the collection of point mutant strains.

### Motility assays

Overnight growth of *C. jejuni* on blood agar plates was harvested and resuspended in phosphate buffered saline to an optical density at 600 nm of 1.0. 0.5 µl aliquots were then injected into semi-solid agar plates (0.4 % w/v agar, 3.7 % w/v brain heart infusion) containing 5×10^−3^ % triphenyl tetrazolium chloride, a redox dye which allows clear visual assessment of growth. The diameter of growth was measured after 16 hours of incubation.

## Data availability

The map for FliD_cj_ is available at EMDB with accession code EMD-10210, and the atomic model is available in Protein Data Bank with accession code 6SIH. The map for the native filament is available at EMDB with accession code EMD-10244. All other data supporting the findings of this study are available from the corresponding authors upon request.

## Acknowledgements

This work was funded by a UK Biotechnology and Biological Sciences Research Council (BBSRC) grant (BB/R009759/1) to J.R.C.B. N.S.A. was recipient of PhD scholarship from The Global Strategic Alliance at the University of Sheffield. A.J.T. was funded by a BBSRC grant (BB/R003491/1) to D.J.K. We thank the members of EM facility for their essential assistance and microscope access, and we acknowledge the members of Prof. Per Bullough’s laboratory for fruitful discussions. Cryo-EM data for FliD_cj_ was collected at the UK national Electron Bio-Imaging centre (eBIC), proposal EM19709-1. The *C.jejuni* filament data was collected at the University of Sheffield Electron Microscopy Facility.

## Author contributions

N.S.A. and J.R.C.B. conceived the project and designed the structural experiments. A.J.T. and D.J.K. designed the *C. jejuni* cloning, mutagenesis and motility assays. N.S.A. performed the protein purification, Cryo-EM data collection and processing, as well as the negative stain experiments. A.J.T. performed the *C. jejuni* mutagenesis and motility assays, together with N.S.A. S.T. provided assistance with data collection and setup of electron microscopy facility. D.F. and F.D. refined the FliDcj atomic model with Rosetta. All authors contributed to the writing and editing of the manuscript.

