## Supplementary figures for "The cryo-EM structure of the bacterial flagellum cap complex suggests a molecular mechanism for filament elongation"

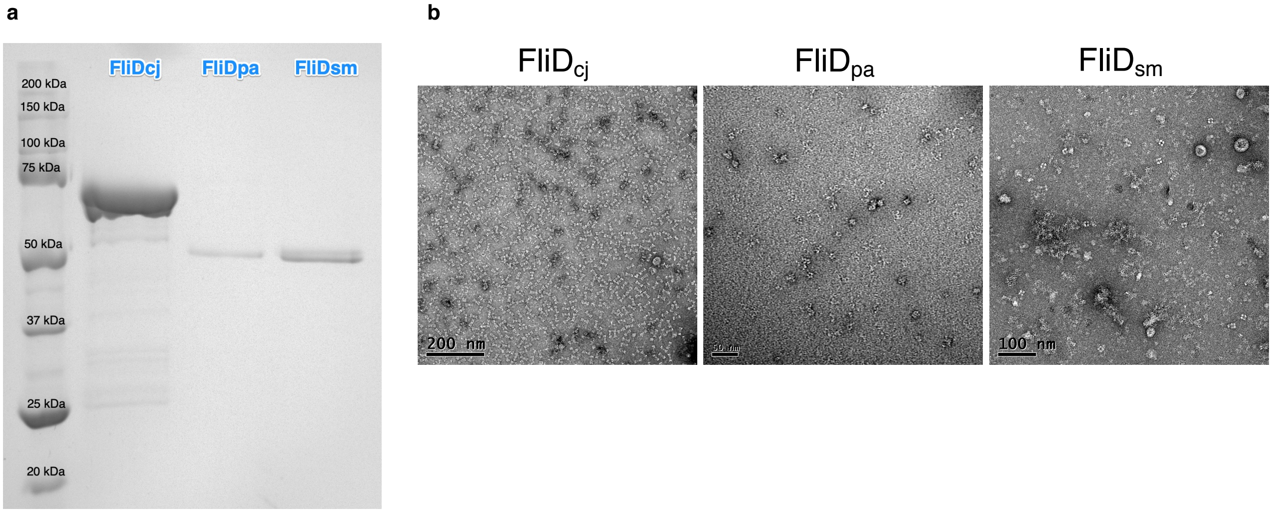


**Supplementary Figure 1: Purification and oligomerization of recombinant FliD. (a)** SDS-PAGE gel showing the purified recombinant $FliD_{cj}$ (70 kDa), $FliD_{pa}$ and $FliD_{sm}$ (50 kDa) proteins. **(b)** Negative stain of the samples in (a), diluted to ~ 0.1 mg/ml. $FliD_{cj}$ appears as well-distributed, homogeneous particles, while $FliD_{pa}$ and $FliD_{sm}$ form heterogeneous particles, with significant aggregation.

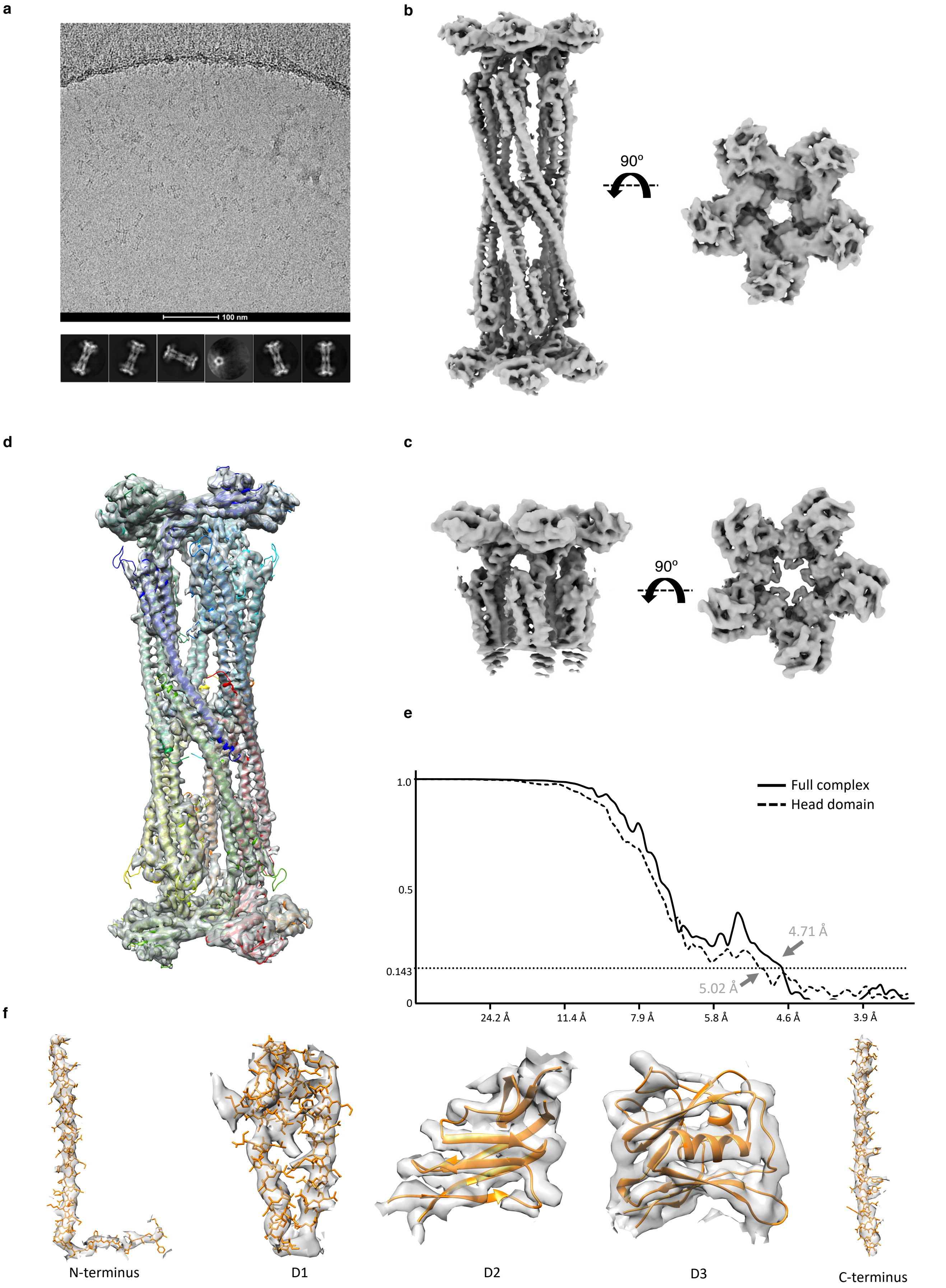


**Supplementary Figure 2: Cryo-EM data for FliD$_{cj}$ flagellum capping protein. (a)** Cryo-electron micrograph of the FliD$_{cj}$ complex. Large particles (~ 30 nm x 5 nm) are visible. Below are 2D classes generated from ~56000 particles. **(b)** Cryo-EM map of the full complex. Side view (left) and top view (right). **(c)** Cryo-EM map of the complex obtained from a masked refinement of (b) allowing for better resolution in the head domains. **(d)** Model of FliDcj built into the density map in (b). **(e)** FSC plots for the maps in (b) and (c) showing the resolution to be 4.71 Å and 5.02 Å respectively. **(f)** Fit of the various regions of FliD$_{cj}$ into the cryo-EM map. The N- and C- termini were manually built into the density in (b) , while D1, D2 and D3 domains were modelled in PHYRE2 (see Materials and Methods).

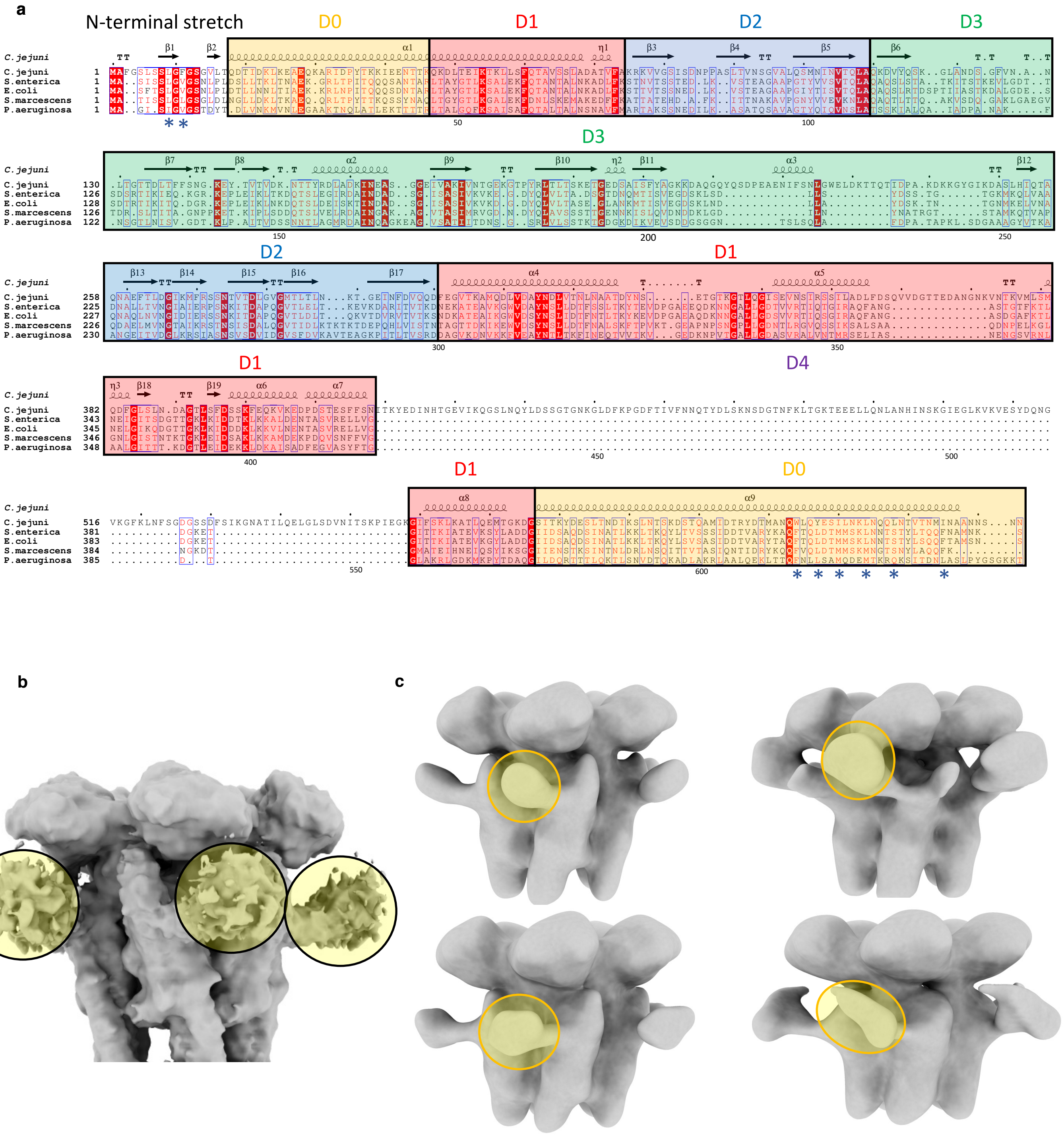


**Supplementary Figure 3: FliD$_{cj}$ specific D4 domain. (a)** Sequence alignment for FliD from different species. The secondary structure elements for FliD$_{cj}$ are shown at the top. The domains are colour coded as follows: *Yellow***:** D0 terminal domains*. Red:* D1 domain. *Blue:* D2 domain. *Green:* D3 domain. *Uncoloured:* D4 domain found only in FliD$_{cj}$. Stars indicate important residues in terminal interactions, for which mutagenesis experiments were performed in Figure 3. **(b)** Low-contour rendering of the head domain-focused cryo-EM map (Figure S2c). Density attributed to the D4 domain is indicated with yellow circles. **(c)** Further 3D classification revealed distinct conformations of the D4 domain, but remained at low resolution because of the low number of particles in each class.

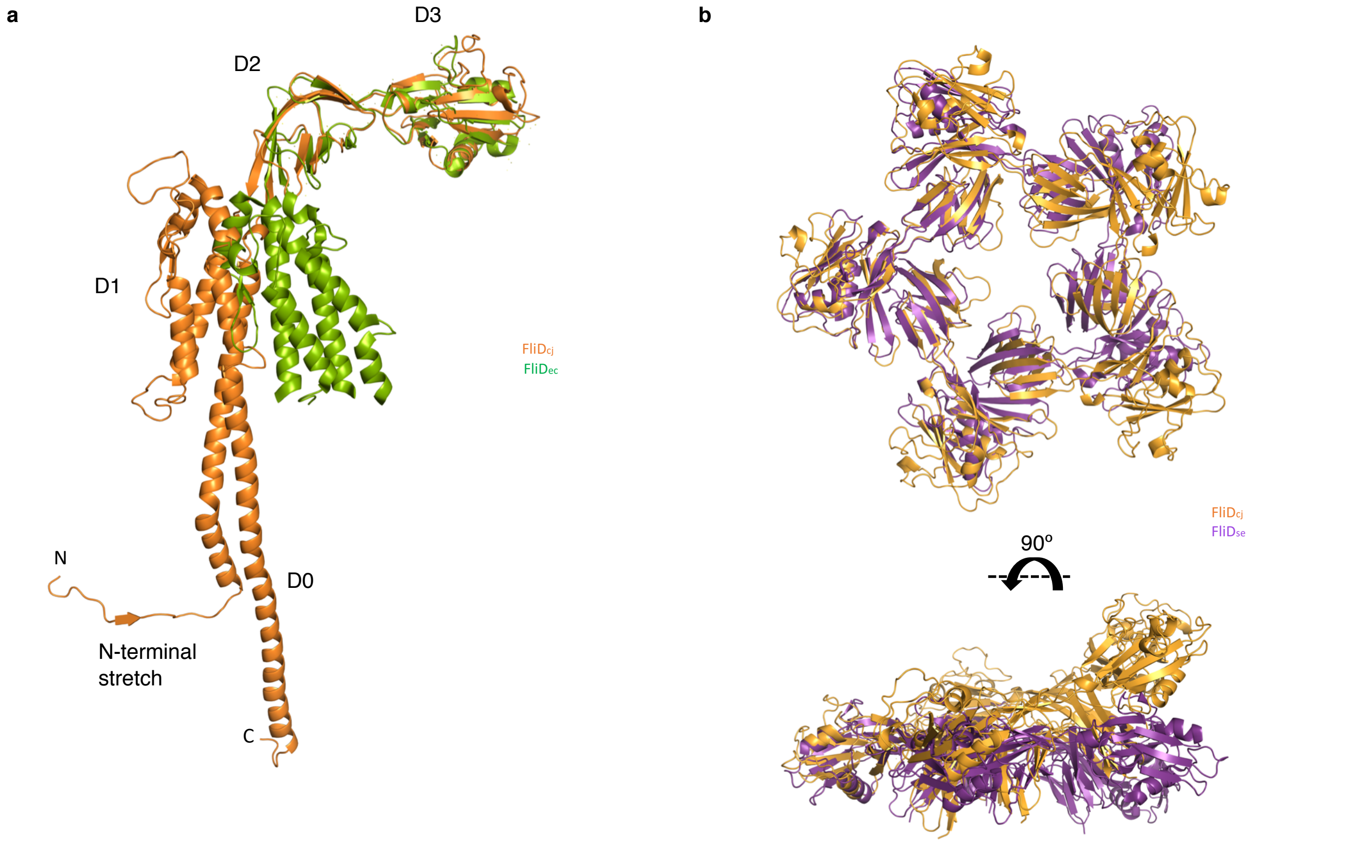


**Supplementary Figure 4 : Comparison of the FliD structures across bacterial species. (a)** Overlay of the FliD structures from *C. jejuni* ($FliD_{cj}$, this study, orange) and *E. coli* ($FliD_{ec}$, 5H5V, green). **(b)** Alignments of X-ray crystallography derived oligomeric structure of the D1 domain from $FliD_{sm}$ to that of the $FliD_{cj}$ pentamer. The angle at which D2-D3 domains are located to D1 in each structure are different.

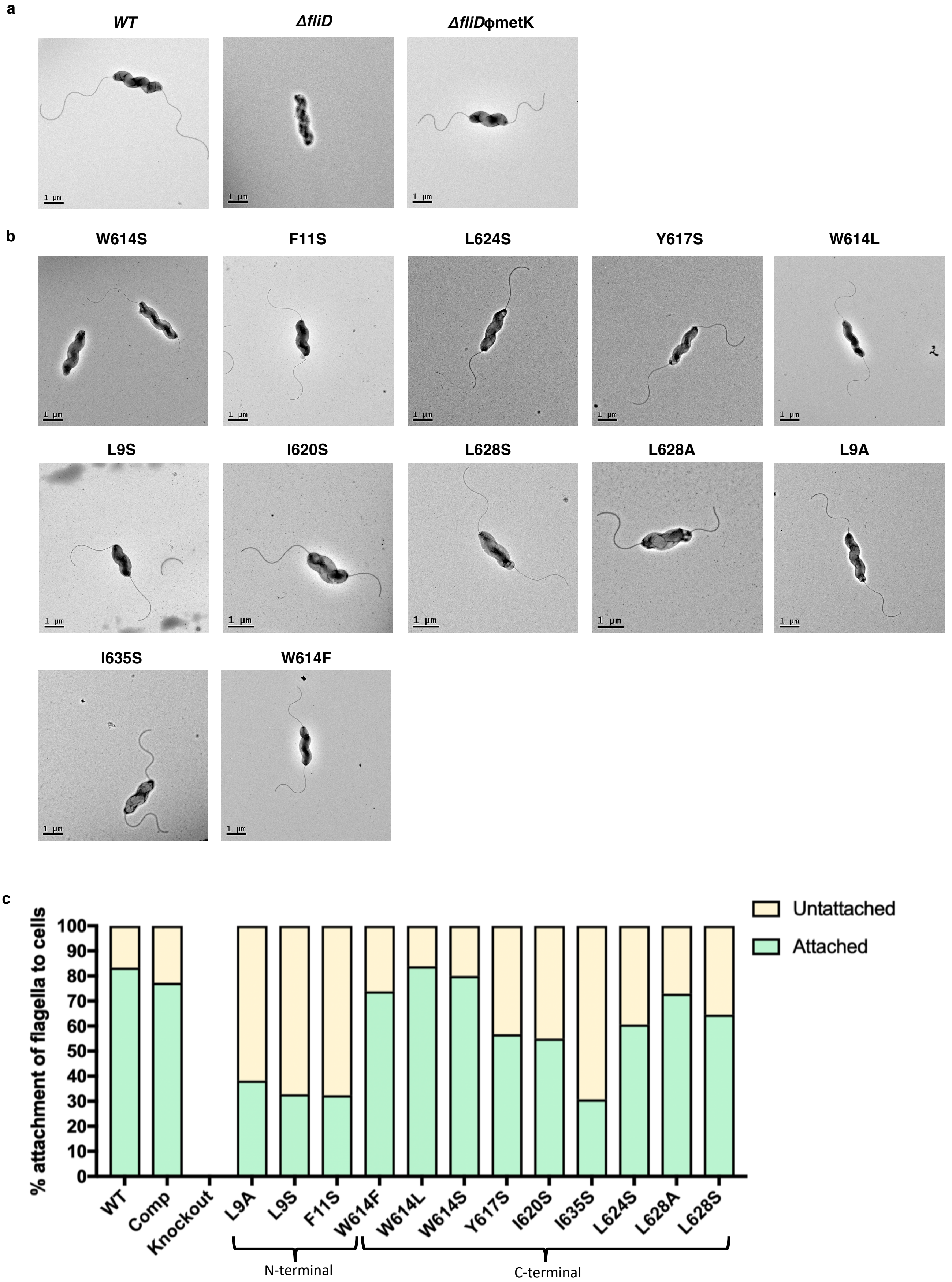


**Supplementary Figure 5: Point mutant motility and flagella attachment. (a)** Negative stain micrographs of cultured *C. jejuni* bacteria from the wild type, *fliD* knockout and complement mutant strains. **(b)** Negative stain micrographs of *C. jejuni* cultures containing FliD point mutants as in in Figure 3. **(c)** Plot of the percentage flagella attachment to cells calculated from micrographs for each point mutant as in Figure 3. Attached are coloured green and unattached yellow..

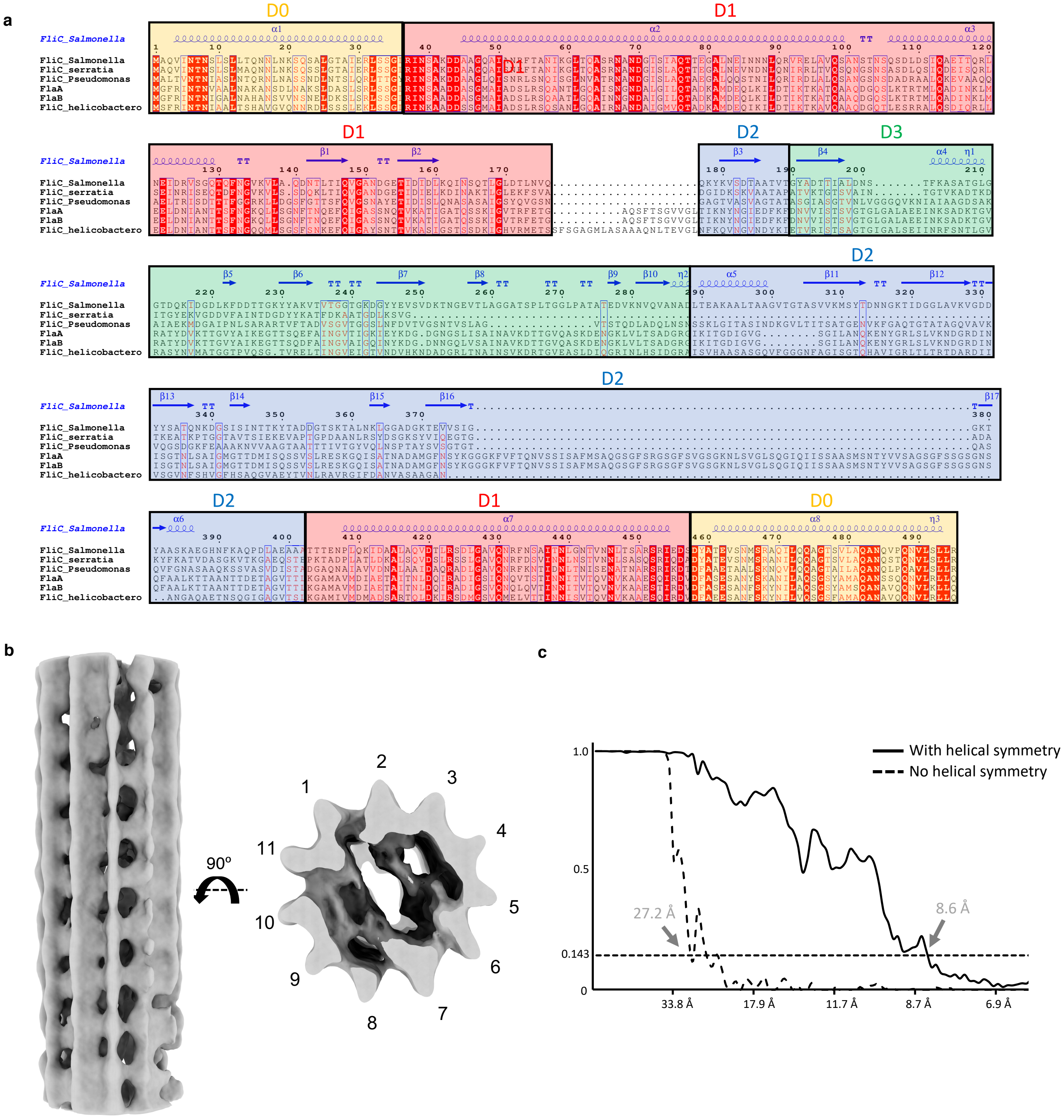


**Supplementary Figure 6: Cryo-EM data for native *C.jejuni* flagella. (a)** Sequence alignment for FliC from different species. The domains are colour coded as follows: *Yellow*: D0 terminal domains. *Red:* D1 domain. *Blue:* D2 domain. *Green:* D3 domain. **(b)** Asymmetric reconstruction of the *C.jejuni* native flagellum filament, with 11 protofilaments visible in the top view. Top view (right) and side view (left). **(c)** FSC curve for the 3D reconstruction of the *C. jejuni* flagellum filament, with (full line) and without (dotted line) helical symmetry.

| Name | Sequence 5' – 3' |
|---|---|
| fliDmutantF1 | GAGCTCGGTACCCGGGGATCCTCTAGAGTCgtcgatataagcttttaactagc |
| fliDmutantR1 | AAGCTGTCAAACATGAGAACCAAGGAGAATgtaatttagttttgatttctgttaa |
| fliDmutantF2 | GAATTGTTTTAGTACCTAGCCAAGGTGTGCatacctctaaagactcaactcag |
| fliDmutantR2 | AGAATACTCAAGCTTGCATGCCTGCAGGTCactgtttcattgttatgcac |
| KanF | ATTCTCCTTGGTTCTCATGTTTGACAGCTTAT |
| KanR | GCACACCTTGGCTAGGTACTAAAACAATTCAT |
| fliDcompF | AATATTCGTCTCACATGgcatttggtagtctatctagttta |
| fliDcompR | AATATTCGTCTCACATGgcttgattttgagaataagc |

**Supplementary Table 1:** Primers for construction of *fliD* deletion mutant and complemented strains. The uppercase sequences of the *fliD* mutant primers are the adaptor regions used in the Gibson assembly cloning, while the lowercase sequences are the regions annealing to a region upstream of fliD (F1) and just inside the *fliD* coding region (R1) or at the end of the *fliD* coding region (F2) and downstream of *fliD* (R2). The KanF and KanR primers are adaptors that also amplify the kan gene from pJMK30

| Name | Sequence 5' – 3' |
|---|---|
| F3L_F | TCATGGCATTaGGTAGTCTATC |
| F3S_F | TCATGGCATcTGGTAGTCTATC |
| F3_R | AAAAGTCCTTTCATTTAAAATGAAC |
| L9A_F | TCTATCTAGTgcAGGATTTGGTTC |
| L9S_F | TCTATCTAGTTcAGGATTTGGTTC |
| L9_R | CTACCAAATGCCATGAAAAAG |
| F11L_F | GTTTAGGATTaGGTTCTGGGG |
| F11S_F | GTTTAGGATcTGGTTCTGGGG |
| F11_R | TAGATAGACTACCAAATGC |
| Y315L_F | GGTGGATGCTctTAATGATTTAGTAAC |
| Y315S_F | GGTGGATGCTTcTAATGATTTAGTAAC |
| Y315_R | AAATCTTGCATGGCTTTTG |
| N316S_F | GATGCTTATAgTGATTTAGTAACCAATC |
| N316L_F | GATGCTTATctTGATTTAGTAACCAATC |
| N316_R | CACCAAATCTTGCATGGC |
| L318A_F | TTATAATGATgcAGTAACCAATCTTAATGC |
| L318S_F | TTATAATGATTcAGTAACCAATCTTAATGC |
| L318_R | GCATCCACCAAATCTTGC |
| L338A_F | AAAAGGAACTgcACAAGGCATC |
| L338S_F | AAAAGGAACTTcACAAGGCATC |
| L338_R | GTTCCAGTTTCACTATTATAG |
| D397N_F | TTTGAGTTTTaATTCTTCTAAATTTGAAC |
| D397L_F | TTTGAGTTTTctTTCTTCTAAATTTGAAC |
| D397_R | GTGCCTGCATCATTTAAAC |
| K400S_F | GATTCTTCTAgtTTTGAACAAAAAGTTAAAGAAGATC |
| K400L_F | GATTCTTCTttATTTGAACAAAAAGTTAAAGAAGATC |
| K400_R | AAAACTCAAAGTGCCTGC |
| L592A_F | TATTAAATCAgcAAATACCTCTAAA |
| L592_R | TCATTTGTCAAACTCTCATC |
| M602L_F | AACTCAGGCTcTGATTGATACAAG |
| M602_R | GAGTCTTTAGAGGTATTTAATG |
| W614L_F | GCGAATCAATtGTTGCAATATG |
| W614S_F | GCGAATCAATcGTTGCAATATG |
| W614F_F | GCGAATCAATtcTTGCAATATG |
| W614_R | CATTGTATCATATCTTGTATCAATC |
| Y617L_F | GGTTGCAATtaGAGAGTATTTTAAATAAAC |
| Y617S_F | GGTTGCAATcTGAGAGTATTTTAAATAAAC |
| Y617_R | ATTGATTCGCCATTGTATC |
| I620A_F | ATATGAGAGTgcTTTAAATAAACTCAATCAACAGC |
| I620S_F | ATATGAGAGTtcTTTAAATAAACTCAATCAACAGC |
| I620_R | TGCAACCATTGATTCGCC |
| L624A_F | TTTAAATAAAgcCAATCAACAGCTAAATACTGTAAC |
| L624S_F | TTTAAATAAAtcCAATCAACAGCTAAATACTGTAAC |
| L624_R | ATACTCTCATATTGCAACC |
| L628A_F | CAATCAACAGgcAAATACTGTAACTAATATG |
| L628S_F | CAATCAACAGtcAAATACTGTAACTAATATG |
| L628_R | AGTTTATTTAAAATACTCTCATATTG |
| I635A_F | AACTAATATGgcTAATGCGGCAAACAATTC |
| I635S_F | AACTAATATGtcTAATGCGGCAAACAATTC |
| I635_R | ACAGTATTTAGCTGTTGATTG |

**Supplementary Table 2:** Primers for construction of *fliD* point mutants in *C.jejuni*. The forward primers (F) contain the mutated base(s) shown in lower case.
